# The genetic variant analyses of SARS-CoV-2 strains; circulating in Bangladesh

**DOI:** 10.1101/2020.07.29.226555

**Authors:** Abu Sayeed Mohmmad Mahmud, Tarannum Taznin, Md. Murshed Hasan Sarkar, Mohammad Samir Uzzaman, Eshrar Osman, Md. Ahasan Habib, Shahina Akter, Tanjina Akhter Banu, Barna Goswami, Iffat Jahan, Md. Saddam Hossain, Md. Salim Khan

## Abstract

Genomic mutation of the virus may impact the viral adaptation to the local environment, their transmission, disease manifestation, and the effectiveness of existing treatment and vaccination. The objectives of this study were to characterize genomic variations, non-synonymous amino acid substitutions, especially in target proteins, mutation events per samples, mutation rate, and overall scenario of coronaviruses across the country. To investigate the genetic diversity, a total of 184 genomes of virus strains sampled from different divisions of Bangladesh with sampling dates between the 10th of May 2020 and the 27^th^ of June 2020 were analyzed. To date, a total of 634 mutations located along the entire genome resulting in non-synonymous 274 amino acid substitutions in 22 different proteins were detected with nucleotide mutation rate estimated to be 23.715 substitutions per year. The highest non-synonymous amino acid substitutions were observed at 48 different positions of the papain-like protease (nsp3). Although no mutations were found in nsp7, nsp9, nsp10, and nsp11, yet orf1ab accounts for 56% of total mutations. Among the structural proteins, the highest non-synonymous amino acid substitution (at 36 positions) observed in spike proteins, in which 9 unique locations were detected relative to the global strains, including 516E>Q in the boundary of the ACE2 binding region. The most dominated variant G614 (95%) based in spike protein is circulating across the country with co-evolving other variants including L323 (94%) in RNA dependent RNA polymerase (RdRp), K203 (82%) and R204 (82%) in nucleocapsid, and F120 (78%) in NSP2. These variants are mostly seen as linked mutations and are part of a haplotype observed in Europe. Data suggest effective containment of clade G strains (4.8%) with sub-clusters GR 82.4%, and GH clade 6.4%.

**Highlights:**

1. We have sequenced 137 and analyzed 184 whole-genomes sequences of SARS-CoV-2 strains from different divisions of Bangladesh.
2. A total of 634 mutation sites across the SARS-CoV-2 genome and 274 non-synonymous amino acid substitutions were detected.
3. The mutation rate of SARS-CoV-2 estimated to be 23.715 nucleotide substitutions per year.
4. Nine unique variants were detected based on non-anonymous amino acid substitutions in spike protein relative to the global SARS-CoV-2 strains.

## 1. Introduction

Most of the coronaviruses are naturally hosted by bats (Cui *et al*., 2019, Li *et al*., 2020), and it is postulated that coronaviruses jumped to humans from the bat reservoir (Dominguez *et al*., 2007, Li *et al*., 2005). Besides bat, coronaviruses also identified in several avian and mammals hosts (Cavanagh, 2007, Ismail *et al*., 2003). Only six coronaviruses have been known to cause infection in humans until late 2019, and these include SARS-CoV, HCoV-OC43, CoV-HKU1, HCoV-NL63, HCoV-229E, and MERS-CoV. A seventh human coronavirus, named SARS-CoV-2, which is closely related to SARS-CoV-1 major cause of pandemic in the modern history of humans.

Based on the whole genome sequencing data, it has been shown that the nucleotides bases of the SARS-CoV-2 virus have 96.2% similarity with that of a bat SARS-related coronavirus (RaTG13) (Paraskevis *et al*., 2020). However, it has a low resemblance to that of MERS-CoV (∼50%) or SARS-CoV (∼79%) (Gralinski and Menachery, 2020, Zhu *et al*., 2020). Coronaviruses mostly associated with mild symptoms (Su *et al*., 2016) with two marked exceptions that caused a major epidemic. The first coronavirus mediated outbreak that caused Severe Acute Respiratory Syndrome Coronavirus (SARS-CoV) occurred in 2002, with infected over 8,000 cases and killed 800 (Graham and Baric, 2010) and not known to circulate since 2003. The other coronavirus that caused Middle East Respiratory Syndrome (MERS-CoV) causing sporadic infections, mostly in the Arabian peninsula in 2012, a less infective but highly lethal virus, which infected 2,294 laboratory-confirmed cases and killed 858 (Cui *et al*., 2019, Graham and Baric, 2010) including 38 deaths in South Korea (Lee *et al*., 2016). As of the 16^th^ of July 2020, the SARS-CoV-2 virus causative agent of COVID-19 has spread to over 216 countries or regions and had caused over 15785641 cases with 640016 deaths worldwide (Who, 2020). The first COVID-19 case was reported in Bangladesh on the 8^th^ of March 2020, and from there on, more than 223453 confirmed cases have been declared with 2928 death (Iedcr, 2020). As COVID-19 is responsible for enormous human casualties and economic loss posing a serious threat to Bangladesh and globally, an understanding of the ongoing situation and the development of strategies to contain the virus’s spread are urgently needed.

Novel beta coronaviruses are positive-sense single-stranded non-segmented enveloped RNA viruses. They belong to some of the largest genomes among RNA viruses, with around 25-32 kb. The genome of circulating novel coronavirus comprises 29903 bases include a (‘-cap, ‘-untranslated region (UTR), ORFs, ‘-UTR, and ‘-poly(A) tail). The ORF 1ab that encodes 16 non-structural proteins (nsps) represents around 71.5 percent of the entire genome, while the remaining ORFs encode structural and accessory proteins. The orf1ab translated to 7097 amino acids (aa) long two viral polyproteins (pp) pp1a and pp1ab by host ribosomes. ORF1a encodes 4406 aa long polyproteins, including two cysteine proteases, a papain-like protease (PL^pro^), and a 3C-like protease (3CL^pro^). PL^pro^ cleaves after glycine residue with recognition site LXGG and generates the first three mature non-structural proteins (nsp), nsp1, 2, and 3. 3CL^pro^ hydrolyzes after amino acid Glutamine residue is responsible for cleavage of the remaining 11 sites resulting in the release of a total of mature 16 non-structural proteins (nsps).

Many nsps assemble to form a replicase-transcriptase complex that is responsible for genomic RNA replication and transcription of the sub-genomic RNAs. The sub-genomic RNA, which is encoded within the ‘ends of the genome serves as mRNAs for the transcription and subsequent translation of the structural and other accessory genes. The primary structural proteins, including envelope protein (E), participate in viral assembly and budding. The membrane protein (M) defines the shape of the viral envelope, nucleocapsid protein (N) binds to the RNA genome and is also involved in viral assembly and budding. Similar to SARS-CoV, spike protein mediates attachment of the virus to the host cell surface receptors Angiotensin-Converting Enzyme 2 (ACE2) and facilitate viral entrance into host cells (Letko *et al*., 2020, Zhou *et al*., 2020). Therefore, the spike protein largely determines host tropism and infectivity of a coronavirus.

Usually, a higher rate of mutation occurred in typical RNA viruses that result in a different version of the viral population with diverse genomes. As SARS-CoV-2 infects human, replicate its copy, and transmits to more individuals they accumulate difference between the progeny and the original viral genome. It has been estimated for coronavirus that the average evolutionary rate is approximately 10^−4^ nucleotide substitutions per site per year (Su *et al*., 2016). When a different variant of viral populations, infecting patients, may contribute to differences in clinical outcomes among the patients. It is, therefore, demand the constant surveillance of the newly arisen viral variant by continuously monitoring the sequences of SARS-CoV-2 from different patients. This study aimed to analyze the variants being circulated in Bangladesh due to mutation at nucleotides and amino acids level.

## 2. Materials and Methods

### 2.1 Novel coronavirus detection and sequencing

The extraction of the viral nucleic acid from the nasopharyngeal specimen was performed using the PureLink™ Viral RNA/DNA Mini Kit (Invitrogen) according to the manufacturer’s protocol. Samples were identified as positive for SARS-CoV-2 (N-gene and ORF 1ab-gene) by Novel Coronavirus (2019-nCoV) Nucleic Acid Diagnostic Kit (Sansure Biotech) according to the manufacturer’s protocol. Random hexamers generated the cDNA directed reverse transcription using 20 μl of RNA extract, 660 μM dNTPs, 5 x RT Improm II reaction buffer (Promega), 50 ng hexanucleotides, 1.5 mM MgCl2, 20 U RNasin® Plus RNase Inhibitor (Promega, Madison, Wisconsin) and 1U of ImProm-II™ Reverse Transcriptase (Promega). SARS-CoV-2 genomes were quantified by using a qRT-PCR assay targeting a conserved region of the envelope gene. Sequencing-ready libraries were prepared using cDNA from the CoV sample (CoVOC43), the viral pool sample (ViralPool) with Nextera Flex for Enrichment (Illumina, Catalog no. 20025524) and IDT for Illumina Nextera DNA UD Indexes (Illumina, Catalog no. 20027213). The total DNA input recommended for tagmentation is 10–1000 ng. After tagmentation and amplification, samples were enrichment with the Respiratory Virus Oligos Panel (Illumina, Catalog no. 20042472), which features ∼7800 probes designed to detect respiratory viruses, recent flu strains, and SARS-CoV-2. After enrichment, the prepared libraries were quantified, pooled, and loaded onto the MiniSeq™ System with an output of 2 × 76-bp paired-end reads for sequencing.

### 2.2 Bioinformatic analysis for generating sequencing data

FASTQ data were exported from the local run manager to BaseSpace Hub, Illumina. DRAGEN RNA Pathogen Detection V3.5.14 generated consensus FASTA. FASTQ data were further analyzed by the IDSeq platform to check the consensus FASTA coverages (Kalantar *et al*., 2020). Consensus FASTA files were uploaded in Genome Detective Virus Tools (Vilsker *et al*., 2018) to check the virus open reading frame and mutations at both nucleotide and amino acids level. The data was post-analyzed by comparing it with the China National Center for Bioinformation (Members and Partners, 2019) and Nextstrain.org.

### 2.3 Dataset construction

We have sequenced a 137 whole-genome of SARS-CoV-2 from COVID-19 patients in Bangladesh. A total of 188 datasets were constructed, including 51 viral sequences from Bangladeshi patients sequenced by other research groups that were submitted to the GISAID database between May 15th and July 10^th^, 2020. Among the isolates 101 from Dhaka, 31 from Chattogram, 22 from Rajshahi, 22 from Rangpur, 9 from Sylhet, and 3 from Khulna division. The reference SARS-CoV-2 sequence, which was isolated in Wuhan, China, downloaded from GeneBank (NC_045512.3)4. Four sequences (EPI_ISL_435057) were excluded from the analysis due to harboring an extreme number of unique mutations, which can result from sequencing errors.

### 2.4 Nucleotide substitution analysis

SARS-CoV-2 isolate sequences from Bangladesh were compared to the reference SARS-CoV-2 sequence (NC_045512.3) using nucleotide substitutions. Sequence alignment for constructed 184 datasets was performed using Multiple Sequence Comparison by Log-Expectation (MUSCLE) software and MEGAX (Kumar *et al*., 2018). Aligned sequences were separately displayed in the AliView to verify that the sequences were in the frame. Degenerated nucleotide was removed, and all the sequences length were maintained equally by removing ‘UTR manually (265 nt) from5’ end and ‘UTR (229 nucleotides) from ‘ends. Nucleotides substitution in both5’ UTR and ‘UTR were separately observed. The amino acid substitution and all nucleotides substitutions were reconfirmed separately by utilizing open-source software Genome detective (Vilsker *et al*., 2018), and China National Center for Bioinformation (CNCB)

## 3. Results

### 3.1 Distribution of sequenced SARS-CoV-2

SARS-CoV-2 isolates were collected from age 8 days to 95 years old COVID-19 patients. The age interval of the patients from Bangladesh was approximately 95. Off the available 169patient’s data, 47.3% were between the age group 20–40, while 34.3 % was between 41-60 and 9.5 % was between 61to 80 (**Fig. 1a**). Viral isolates were collected from 64% of male patients and 36% of females (**Fig. 1b**). Most of the sequenced samples come from the capital city Dhaka 101, followed by 31 from Chattogram (**Fig. 1c**). Interestingly, 82.4% of all sequenced SARS-CoV-2 in Bangladesh belong to the GISAID GR clade (**Table 1**), followed by a 6.4% GH clade (**Fig. 1c**). We also observed 9 SARS-CoV-2 belongs to G clade, 5, and 6 cases of coronaviruses were S and O clades, respectively. Although no V clade variants found, one L clade coronavirus (EPI_ISL_458133) was observed. Most of the coronaviruses were pangolin lineage B.1.1 (84%), followed by 7.4% B.1.36 (**Fig. 1d**).

**Table 1:** Categorization of Bangladeshi SARS-CoV-2 strains according to the GISAID clade and pangolin lineages.

| **Clade** | **Nucleotide Changes** | **Amino acid changes** | **No. of variants** |
| --- | --- | --- | --- |
| **G**<br>**(B.1/B.1.1/**<br>**B.1.36)** | 241C>T, 3037C>T<br>14408C>T, 23403A>G | 5'UTR, NSP3:106F<br>NSP12b: 314P>L, S: 614D>G | 9 |
| **GH**<br>**(B.1/B.1.36)** | 241C>T, 3037C>T<br>14408C>T, 23403A>G 25563G>T | 5'UTR, NSP3:106F<br>NSP12b: 314P>L, S: 614D>G<br>ORF3a: 57Q>H | 12 |
| **GR**<br>**(B.1.1)** | 241C>T, 3037C>T<br>14408C>T, 23403A>G 28881<br>GGG>AAC | 5'UTR, NSP3:106F<br>NSP12b: 314P>L, S: 614D>G<br>N: 203 RG >KR | 155 |
| **V (B2)** | 11083G>T, 26144G>T | NSP6:37L>F, ORF3a: 251G>V | 0 |
| **S (A)** | 8782C>T, 28144T>C | NSP4:76S, ORF8:84L>S | 5 |
| **L** | WIV04-reference sequence | 0 | 1 |
| **O** | Others |  | 6 |

**Fig.1:**
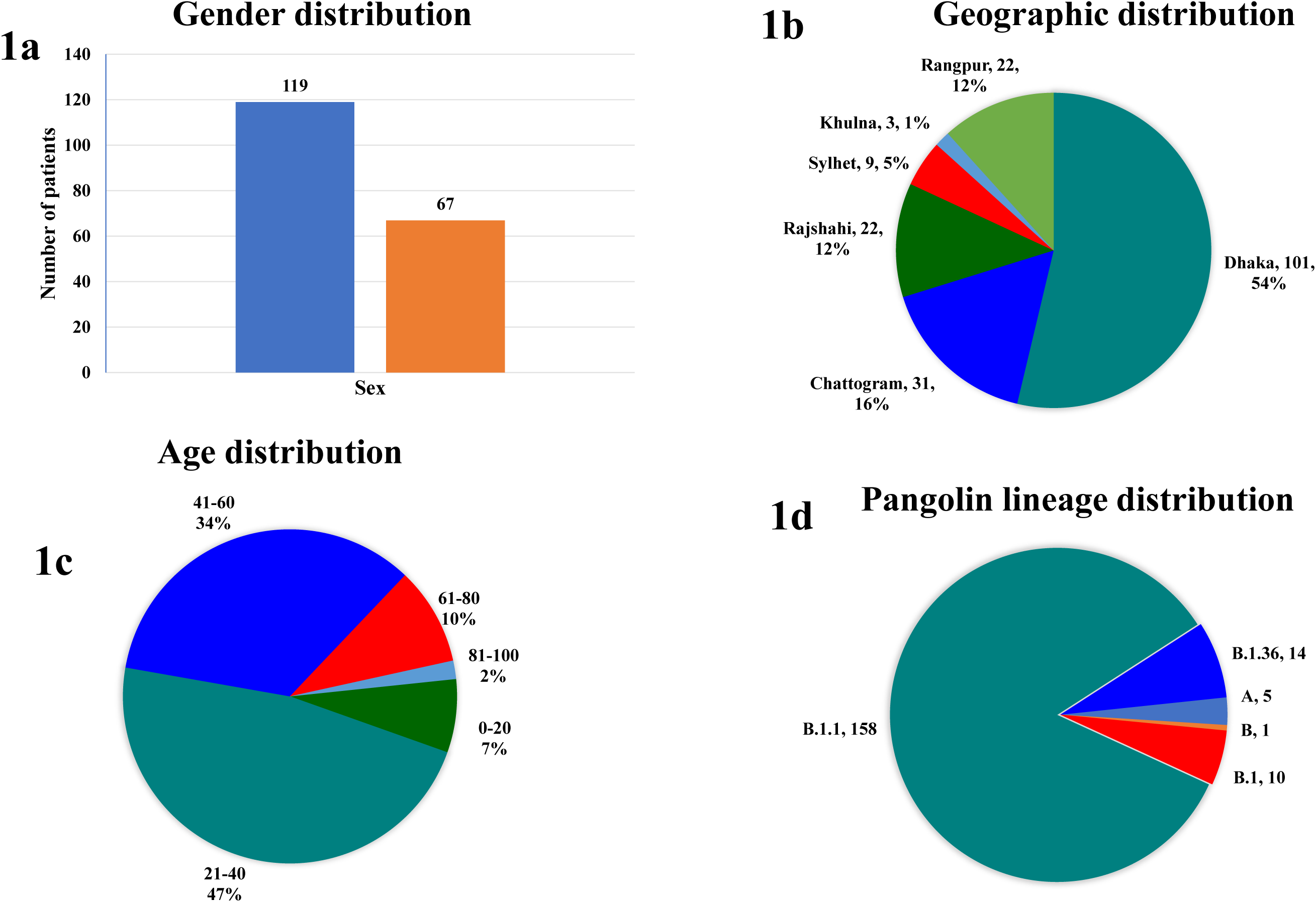
Distribution of sequenced SARS-CoV-2 based on the gender (1a), geography (1b), GISAID clade (1c), and Pangolin lineage (1d).

### 3.2 Variant analysis

Analysis of 184 SARS-CoV-2 highlights a total of 2242 nucleotide mutation events compared to the reference genome (NC_045512.3), with an average of 12.2 mutation events per sample. At the amino acid level, 1258 mutation events were observed with an average of 6.8 per sample. However, these mutations have occurred at a total of 631 different positions of 22 different proteins. Most of the mutations were SNP 575, followed by 43 deletions, 9 insertional, and 4 indels (**Fig 2a**). Of the 575 SNP mutations, thymine replaces cytosine, guanine, and adenine 376 times. At the same time, thymine replaced by cytosine, adenine, and guanines all together by only 65 times (**Fig. 2b**). The highest mutation was observed in the nsp3 or papain protease; it undergoes 48 amino acid mutations (**Fig.3**). Although orf1ab covered 71.3 % of the entire coronavirus genome, only 55.5% of active mutations were in the translated proteins of orf1ab. There is no mutation observed in four non-structural proteins, including nsp7, nsp9, nsp10, and nsp11.

**Fig.2:**
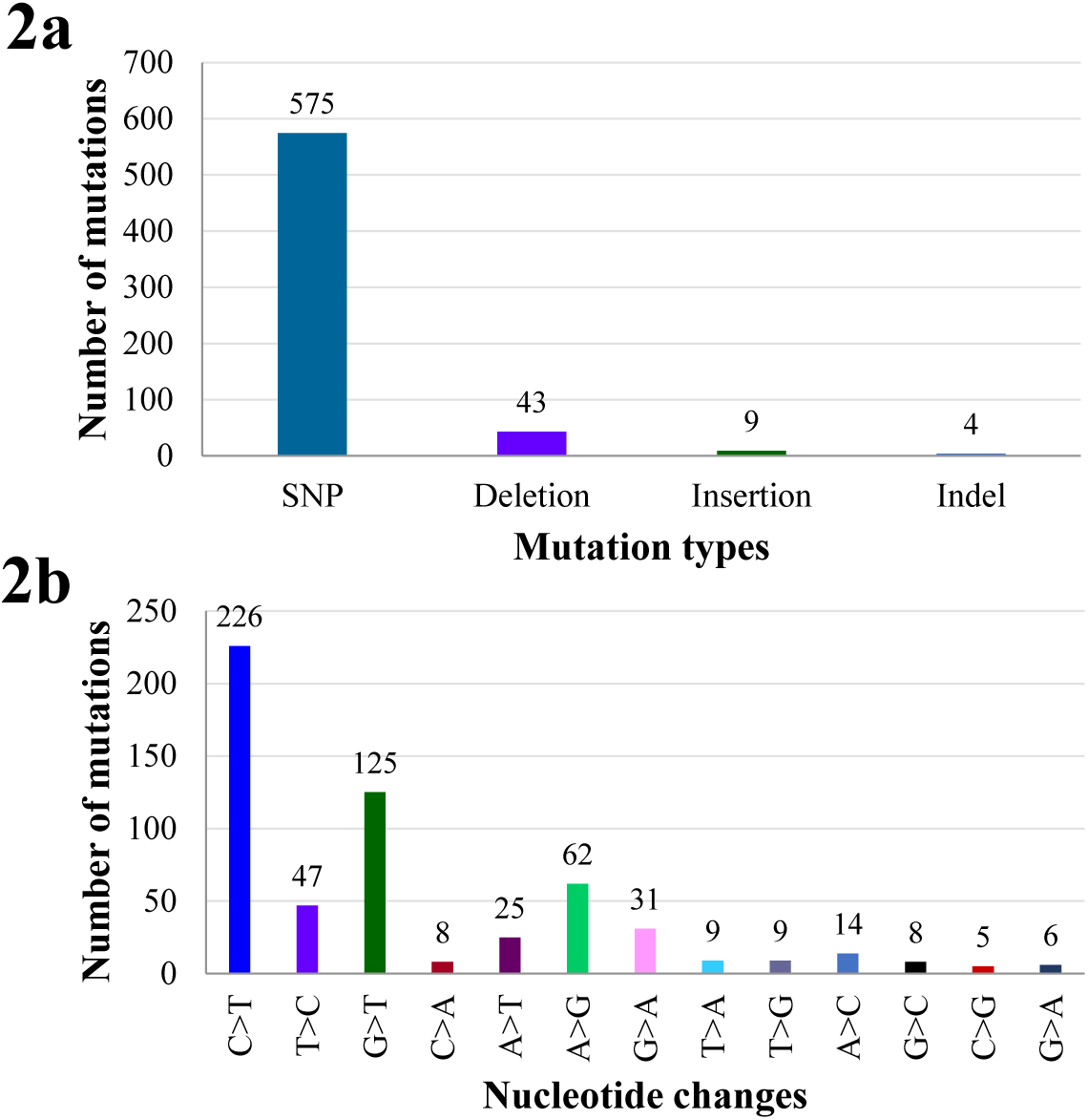
Different mutations type (2a) and the distribution of bases due to SNP (2b).

Papain like protease (PL^pro^) is 1945 amino acid long, one of two proteases of SARS-CoV-2, which generates the first three mature non-structural proteins. The highest number of mutations occurred at 71 positions of PL^pro^ encoding genome, rendering 48 amino acid change. Whereas the main virulence factors 3 like cysteine (3CL^pro^) undergoes less mutational changes, nucleotides have changed at 12 positions resulting in an alteration of 8 amino acids. Another vital enzyme RNA dependent RNA polymerase (RdRp) also undergoes 12 mutations due to nucleotides change at 27 positions. The most recurrent variant based on variations in the RdRp was 14408C>T, followed by 15324C>T with a mutation rate of 95% and 4.3%, respectively. Mutation at 14408C>T positions of RdRp renders the change of amino acid proline to Leucine.

The second highest mutations were observed in the S protein after PL^pro^. Mutations were found at 60 different positions of spike proteins, 61% of those mutations rendered the amino acid changes at 36 different positions. The most dominant variant G614 (23403A>G) found 96% of 184 cases due to mutation at position 23403A>G. The second highest variants due to nucleotide changes at position 22444C>T of spike protein covered 6% of cases. The unique 9 variants of SARS-CoV-2 based on a mutation in spike proteins have been found in Bangladesh. These variants include 13S>I, 14Q>H, 26P>L, 140F del, 144Y del, 211N>Y, 516E>Q, 594G>S, and 660Y>F. The change of amino acids glutamic acid at 516 positions to glutamine occurred within the receptor-binding domain (**Fig. 3**). All the mutations were highlighted in figure, also highlighted the mutations at 516, and 518 positions of spike proteins (**Fig. 4**). Another critical variant has arisen in Bangladesh due to changes of amino acids glycine to acidic serine at position 594 of spike protein S1 subunit (**Table 2**). G614 variants were found in 96% of cases, 60 mutations were observed in 98 positions of the spike protein in out of 186 sequenced SARS-CoV-2; however, 36 mutations (61%) have rendered the amino acid changes in the spike protein.

**Table 2:**
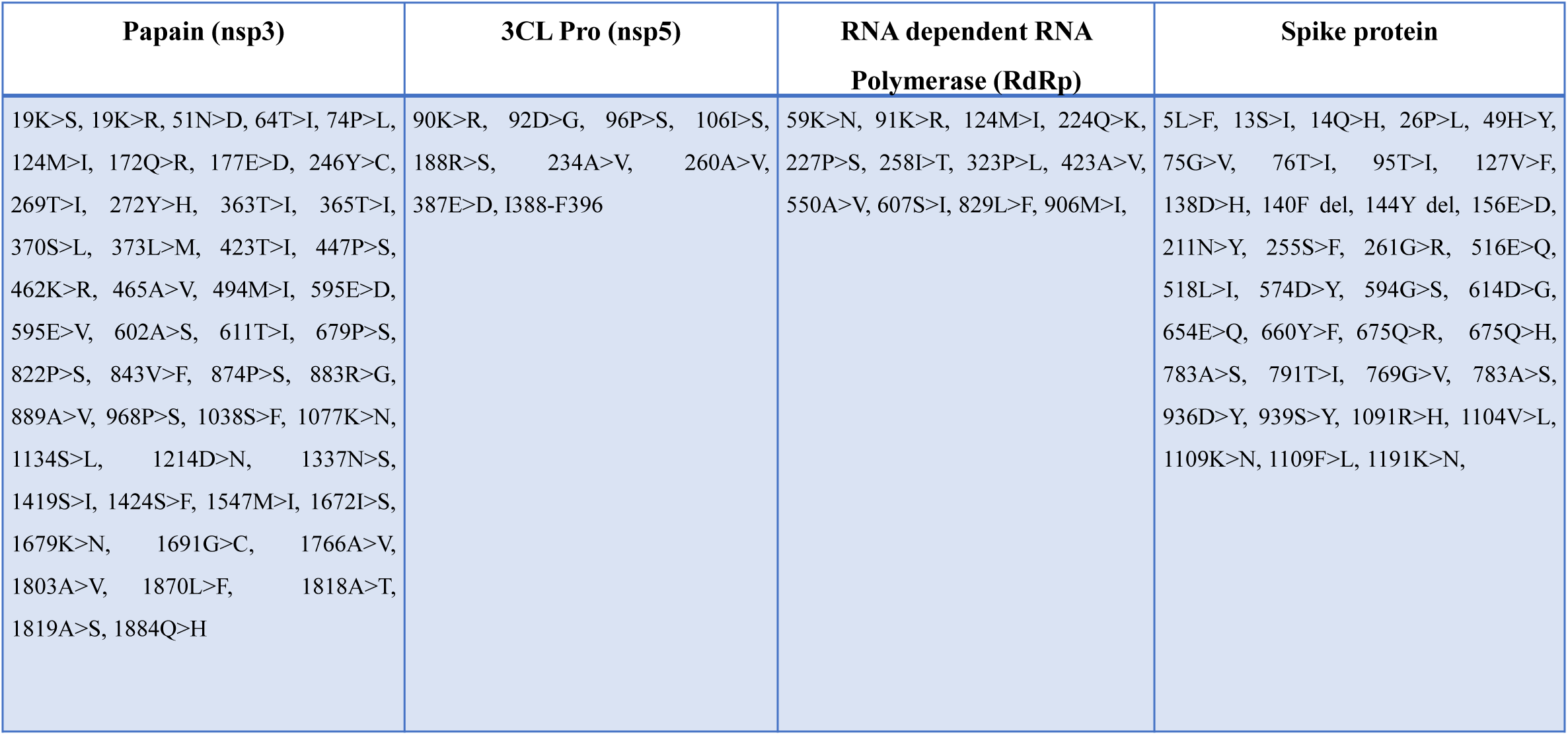
List of detected non-synonymous amino acid substitutions in major target proteins

| Papain (nsp3) | 3CL Pro (nsp5) | RNA dependent RNA<br>Polymerase (RdRp) | Spike protein |
| --- | --- | --- | --- |
| 19K>S, 19K>R, 51N>D, 64T>I, 74P>L,<br>124M>I, 172Q>R, 177E>D, 246Y>C,<br>269T>I, 272Y>H, 363T>I, 365T>I,<br>370S>L, 373L>M, 423T>I, 447P>S,<br>462K>R, 465A>V, 494M>I, 595E>D,<br>595E>V, 602A>S, 611T>I, 679P>S,<br>822P>S, 843V>F, 874P>S, 883R>G,<br>889A>V, 968P>S, 1038S>F, 1077K>N,<br>1134S>L, 1214D>N, 1337N>S,<br>1419S>I, 1424S>F, 1547M>I, 1672I>S,<br>1679K>N, 1691G>C, 1766A>V,<br>1803A>V, 1870L>F, 1818A>T,<br>1819A>S, 1884Q>H | 90K>R, 92D>G, 96P>S, 106I>S,<br>188R>S, 234A>V, 260A>V,<br>387E>D, I388-F396 | 59K>N, 91K>R, 124M>I, 224Q>K,<br>227P>S, 258I>T, 323P>L, 423A>V,<br>550A>V, 607S>I, 829L>F, 906M>I, | 5L>F, 13S>I, 14Q>H, 26P>L, 49H>Y,<br>75G>V, 76T>I, 95T>I, 127V>F,<br>138D>H, 140F del, 144Y del, 156E>D,<br>211N>Y, 255S>F, 261G>R, 516E>Q,<br>518L>I, 574D>Y, 594G>S, 614D>G,<br>654E>Q, 660Y>F, 675Q>R, 675Q>H,<br>783A>S, 791T>I, 769G>V, 783A>S,<br>936D>Y, 939S>Y, 1091R>H, 1104V>L,<br>1109K>N, 1109F>L, 1191K>N, |

**Fig.3:**
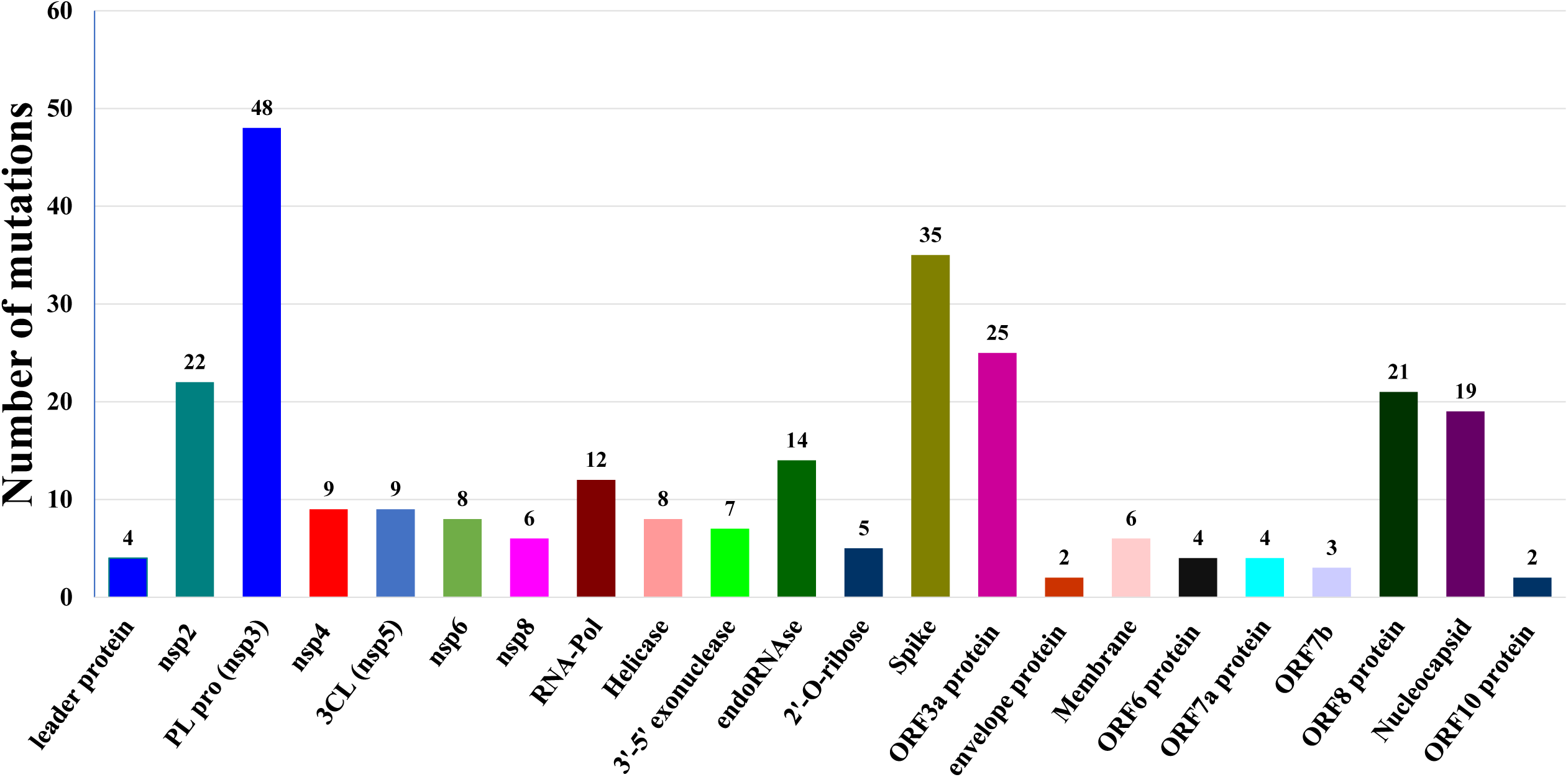
Non-synonymous amino acids substitution in 12 non-structural proteins, 4 structural proteins, and 6 accessory proteins.

**Fig. 4:**
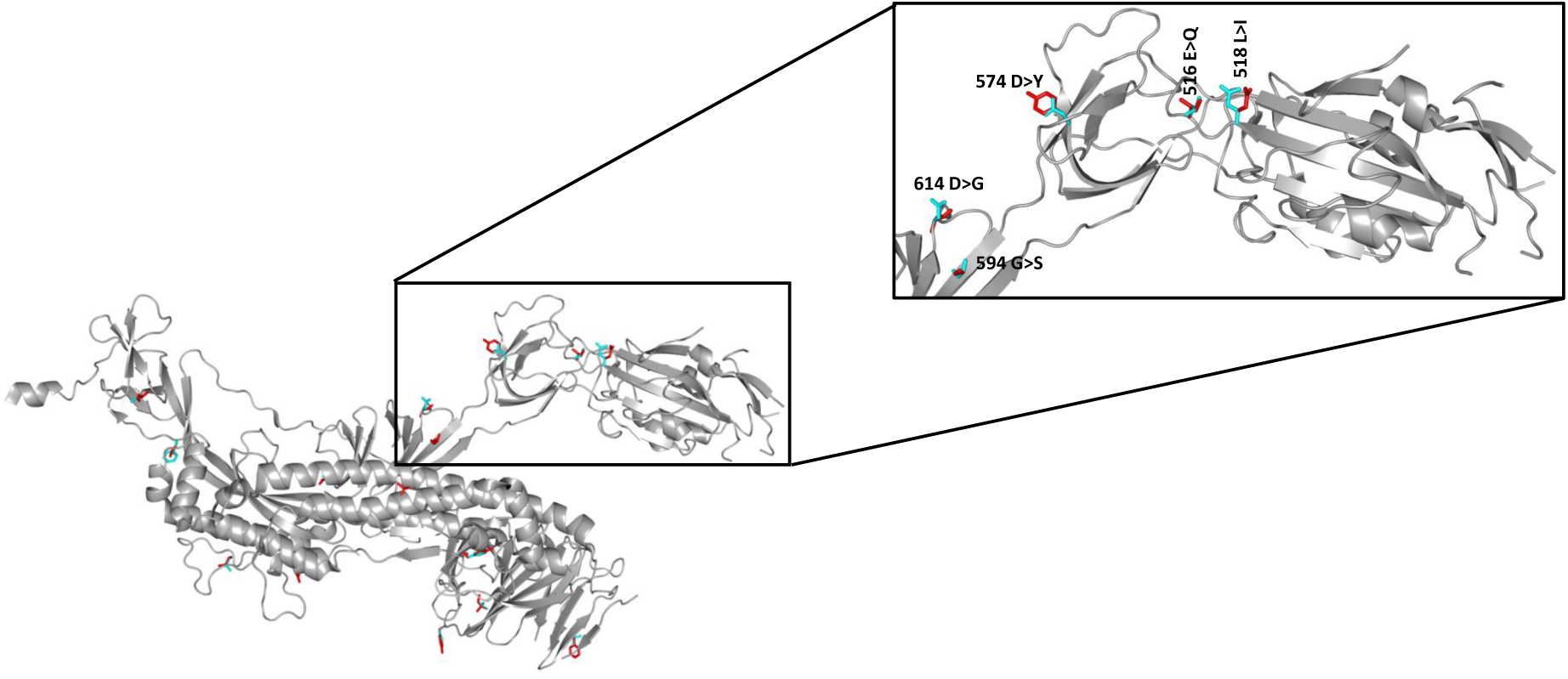
Mutated amino acids are modeled in spike protein structure (PDB ID: 6vxx) using COOT software (Emsley et al., 2010). The structure figures were produced using PYMOL (2). Spike proteins are shown in cartoon representation, mutated amino acids and their corresponding original residues are shown in stick representation in green and magenta color, respectively. Residues highlighted in the box are involved in ACE2 binding.

Nucleotides cytosine has changed into uracil at 241 of 5’UTR found in 95% of cases. Mutation in the nucleotide of nsp2 at 1163 position has changed the Isoleucine to Phenylalanine in 78.3% of cases. In nsp3, mutation at position 3037 found in 94.6% of cases; however, this mutation did not change the amino acid. We also observed nucleotide cytosine at 14408 changed to thymine (uracil), which has altered amino acid proline to Leucine at 323 positions of RNA dependent RNA polymerase. This mutation was observed in 96.7% of cases. The Arg residue at 203 positions of nucleocapsid has changed into Lys residue in 82.6 of cases, while Gly residue at 204 of nucleocapsid changed to Arg residue in 79% of cases. 3CLpro cuts the polyproteins translated from viral RNA to yield functional viral proteins. One of the best-characterized drug targets among coronaviruses is the main protease (Mpro, also called 3CLpro) (Anand *et al*., 2003). Inhibiting the activity of this enzyme would block viral replication. Because no human proteases with a similar cleavage specificity are known, such inhibitors are unlikely to be toxic.

## 4. Discussion

The first Bangladeshi cases of COVID-19 caused by SARS-Cov-2 were reported on the 8^th^ of March 2020, and so far, above 2500 deaths have been documented due to this pandemic. We have analyzed 184 out of 188 SARS-CoV-2 whole-genome sequences available in GISAID published from Bangladesh dated 10^th^ of July 2020. Off the 188 SARS-CoV-2 whole-genome sequenced published to date, four whole-genome sequences (EPI_ISL_445217, EPI_ISL_468071, EPI_ISL_468072, and EPI_ISL_477140) were found very low in quality hence not genetically analyzed in this study. Off the 184 samples examined, variant G614 based on a mutation in spike proteins found in 178 samples, which determines the virus clades G along with frequently co-evolving variants L323, were also found in 178 samples. Other mutations found in 152 samples were K203, and R204 based on mutations in the nucleocapsids of the virus. Overall, 152 sequenced SARS-CoV-2 belong to the GR clade, 12 GH clade, and 9 G clade indicating the viruses are mostly linked to the European origin. The variants C241T located in the non-coding 5’UTR’ were found in 175 samples. Mutation at 3037 positions of the genome found in 174 samples where cytosine changed to thymine located in the nsp3 or M^pro^ did not change amino acids. Other most frequent variants were C14408T located in the RNA dependent RNA polymerase rendered the amino acid Pro to Leu found in 178 samples. These mutations have been detected across Europe, which mostly belongs to GISAID clade G. This viral journey from Wuhan to Bangladesh, lasting 5 months, is documented by 12.2 mutations events per sample, whereas a common mutation per sample worldwide is 7.23 (Mercatelli and Giorgi, 2020).

As more genomes from Bangladesh are made publicly available, analysis of genetic variants has revealed the highest diversity occurring in nsp3 along with spike proteins, nucleocapsids, ORF3a, and ORF8 across the samples. The primary determinant of the host range and pathogenicity of SARS-CoV-2 is its surface-associated S protein. Spike is a trimeric glycoprotein composed of three chains, each chain consisting of two subunits. The S1 subunit located at N-terminal while the S2 subunit located at the C-terminal. The S1 subunit is responsible for recognizing and binding to the host cell receptor ACE2 while the S2 subunit is specializing for membrane fusion. Compared with the S1, the S2 subunit shows much lower variability (Masters, 2006). Although 1721 mutation sites are identified so far in S protein, only 906 lead to amino acid changes. Amino acids 336 to 516 of 1274 amino acid long spike protein form the binding domain, which mediates coronavirus attachment with ACE2 receptors. It has been found that 111 changed amino acids located in the Receptor Binding Domain (RBD). Among them, 8 out of 17 critical amino acids are responsible for protein interactions. As of 12^th^ of July 2020, a total of 60 mutation sites were identified in the spike protein, of which 36 lead to amino acid changes. The most common mutations which dominated and spread all over Bangladesh were G614 found in 178 samples, followed by L26 and F5 found in 5 and 4 samples, respectively.

G614 variant of SARS-CoV-2 isolates predominates over time in locales indicating that this change in the spike protein enhances viral transmission. In a study, it has been shown that retroviruses pseudotyped with SG614 can infect angiotensin-converting enzyme expressing cells significantly more effective way than the SD614 (Zhu *et al*., 2020). However, it has been shown in a separate study that lower RT-qPCR CT value, which is indicative of higher titer of coronavirus variant G614 linked to higher viral load at the upper respiratory tract and thus attenuated disease severity (Korber *et al*., 2020).

Nucleocapsid (N) protein is encoded by a highly variable region of the genes N gene, responsible for the formation of the helical nucleocapsid. N protein can elicit both cell-mediated and humoral immune response and thus has potential value in vaccine development (Ding *et al*., 2016). In the N gene, one nucleotide change was detected at position G2881A found in 152 samples, and two nucleotides changes in the adjacent codons found in 149 samples. These changes have led to alter the two amino acids replacing Arg to Lys, and Gly to Arg at position 203 and 204, respectively. However, the mutations, as mentioned above, did not yet to be associated with changes in viral transmissibility or pathogenicity.

It has suggested based on selection analysis that as the virus is transmitted between humans, the genes of both spike and nucleocapsid undergo episodic selection (Benvenuto *et al*., 2020). The positive selection for parts of any emerging virus is typical (Sironi *et al*., 2015). Mutations in the spike or nucleocapsids genes, and subsequently, their adaptation could affect virus stability and pathogenicity (Baric *et al*., 1997).

SARS-CoV-2 has eight accessory proteins including orf3a, orf3b, orf6, orf7a, orf7b, orf8, orf9b, and orf14 (Ceraolo and Giorgi, 2020). Although ORF14 was not detected in 184 sequencing data analyzed in this study. Among the accessory proteins, ORF3a contains six functional domains (I to VI), which may link to viral infectivity, virulence, and ion channel formation (Issa *et al*., 2020). In a previous study based on 2782 whole-genome sequencing data, it has shown that 51 different non-synonymous amino acid substitutions in the 3a proteins (Issa *et al*., 2020). Whereas, non-synonymous 25 amino acid substitutions were detected in ORF3a in this study. Besides, 21 non-synonymous mutations were detected in ORF8.

In conclusion, our analysis sheds light on the different variants of SARS-CoV-2 currently circulating in Bangladesh. Most recurrent synonymous mutations were detected at positions 241 in 5’UTR, and 3037 in nsp3. Whereas non-synonymous recurrent mutations were detected at positions 1063, 14408, 23403, 29881-29883 in nsp2, nsp12, Spike protein, and nucleocapsid, respectively. These variants indicate the underlying link of Bangladeshi SARS-CoV-2 isolates with part of a haplotype observed high in Europe. Further research should be done to monitor the genetic and non-anonymous variants circulating in Bangladesh for understanding the infectivity and transmission of SARS-CoV-2.

## Acknowledgments

The authors acknowledge the Bangladesh Council of Scientific and Industrial Research for funding the study, submitters of SARS-CoV-2 sequence data to the GISAID database, the database managers, developers, and scientists associated with GISAID.

## References

Anand, K., Ziebuhr, J., Wadhwani, P., Mesters, J. R. & Hilgenfeld, R. (2003). Coronavirus main proteinase (3CLpro) structure: basis for design of anti-SARS drugs. Science, 300(5626) 1763.

Baric, R. S., Yount, B., Hensley, L., Peel, S. A. & Chen, W. (1997). Episodic evolution mediates interspecies transfer of a murine coronavirus. Journal of Virology, 71 (3) 1946–1955.

Benvenuto, D., Giovanetti, M., Ciccozzi, A., Spoto, S., Angeletti, S. & Ciccozzi, M. (2020). The 2019-new coronavirus epidemic: Evidence for virus evolution. J Med Virol, 92 (4) 455–459.

Cavanagh, D. (2007). Coronavirus avian infectious bronchitis virus. Veterinary Research, 38 (2) 281–297.

Ceraolo, C. & Giorgi, F. M. (2020). Genomic variance of the 2019-nCoV coronavirus. J Med Virol, 92 (5) 522–528.

Cui, J., Li, F. & Shi, Z.-L. (2019). Origin and evolution of pathogenic coronaviruses. Nature Reviews Microbiology, 17 (3) 181–192.

Ding, B., Qin, Y. & Chen, M. (2016). Nucleocapsid proteins: roles beyond viral RNA packaging. Wiley interdisciplinary reviews RNA, 7 (2) 213–226.

Dominguez, S. R., O’shea, T. J., Oko, L. M. & Holmes, K. V. (2007). Detection of group 1 coronaviruses in bats in North America. Emerging Infectious Diseases, 13(9) 1295.

Graham, R. L. & Baric, R. S. (2010). Recombination, Reservoirs, and the Modular Spike: Mechanisms of Coronavirus Cross-Species Transmission. Journal of Virology, 84 (7) 3134–3146.

Gralinski, L. E. & Menachery, V. D. (2020). Return of the Coronavirus: 2019-nCoV. Viruses, 12 (2) 135.

Iedcr (2020). Updates on the coronavirus disease 2019 (covid-19) situation in Bangladesh. https://www.iedcr.gov.bd/.

Ismail, M. M., Tang, A. Y. & Saif, Y. M. (2003). Pathogenicity of turkey coronavirus in turkeys and chickens. Avian diseases, 47 (3) 515–522.

Issa, E., Merhi, G., Panossian, B., Salloum, T. & Tokajian, S. (2020). SARS-CoV-2 and ORF3a: Non-synonymous Mutations, Functional Domains, and Viral Pathogenesis. mSystems, 5 (3) e00266–00220.

Kalantar, K., Carvalho, T., De Bourcy, C. F. A., Dimitrov, B., Dingle, G., Egger, R., Han, J., Holmes, O., Juan, Y.-F., King, R., Kislyuk, A., Mariano, M. Reynoso, Cruz, D. R., Sheu, J., Tang, J., Wang, J., Zhang, M., Zhong, E., Ahyong, V., Lay, S., Chea, S., Bohl, J., Manning, J., Tato, C. & Derisi, J. (2020). IDseq –An Open Source Cloud-based Pipeline and Analysis Service for Metagenomic Pathogen Detection and Monitoring. bioRxiv.

Korber, B., Fischer, W. M., Gnanakaran, S., Yoon, H., Theiler, J., Abfalterer, W., Hengartner, N., Giorgi, E. E., Bhattacharya, T., Foley, B., Hastie, K. M., Parker, D., Partridge, D. G., Evans, C. M., Freeman, T. M., De Silva, T. I., Mcdanal, C., Perez, L. G., Tang, H., Moon-Walker, A., Whelan, S. P., Labranche, C. C., Saphire, E. O., Montefiori, D. C., Angyal, A., Brown, R. L., Carrilero, L., Green, L. R., Groves, D. C., Johnson, K. J., Keeley, A. J., Lindsey, B. B., Parsons, P. J., Raza, M., Rowland-Jones, S., Smith, N., Tucker, R. M., Wang, D. & Wyles, M. D. (2020). Tracking changes in SARS-CoV-2 Spike: evidence that D614G increases infectivity of the COVID-19 virus. Cell.

Kumar, S., Stecher, G., Li, M., Knyaz, C. & Tamura, K. (2018). MEGA X: Molecular Evolutionary Genetics Analysis across Computing Platforms. Molecular Biology and Evolution, 35 (6) 1547–1549.

Lee, J., Chowell, G. & Jung, E. (2016). A dynamic compartmental model for the Middle East respiratory syndrome outbreak in the Republic of Korea: A retrospective analysis on control interventions and superspreading events. Journal of Theoretical Biology, 408 118–126.

Letko, M., Marzi, A. & Munster, V. (2020). Functional assessment of cell entry and receptor usage for SARS-CoV-2 and other lineage B betacoronaviruses. Nature Microbiology, 5 (4) 562–569.

Li, W., Shi, Z., Yu, M., Ren, W., Smith, C., Epstein, J. H., Wang, H., Crameri, G., Hu, Z. & Zhang, H. (2005). Bats are natural reservoirs of SARS-like coronaviruses. Science, 310 (5748) 676–679.

Li, X., Song, Y., Wong, G. & Cui, J. (2020). Bat origin of a new human coronavirus: there and back again. Science China Life Sciences, 63 (3) 461–462.

Masters, P. S. (2006). The molecular biology of coronaviruses. Advances in Virus Research, 66 193–292.

Members, N. G. D. C. & Partners (2019). Database Resources of the National Genomics Data Center in 2020. Nucleic Acids Research, 48 (D1) D24–D33.

Mercatelli, D. & Giorgi, F. M. (2020). Geographic and Genomic Distribution of SARS-CoV-2 Mutations. Frontiers in Microbiology, 11 (1800).

Paraskevis, D., Kostaki, E. G., Magiorkinis, G., Panayiotakopoulos, G., Sourvinos, G. & Tsiodras, S. (2020). Full-genome evolutionary analysis of the novel corona virus (2019-nCoV) rejects the hypothesis of emergence as a result of a recent recombination event. Infection, Genetics and Evolution, 79 104212.

Sironi, M., Cagliani, R., Forni, D. & Clerici, M. (2015). Evolutionary insights into host-pathogen interactions from mammalian sequence data. Nature Reviews Genetics, 16 (4) 224–236.

Su, S., Wong, G., Shi, W., Liu, J., Lai, A. C. K., Zhou, J., Liu, W., Bi, Y. & Gao, G. F. (2016). Epidemiology, Genetic Recombination, and Pathogenesis of Coronaviruses. Trends in microbiology, 24 (6) 490–502.

Vilsker, M., Moosa, Y., Nooij, S., Fonseca, V., Ghysens, Y., Dumon, K., Pauwels, R., Alcantara, L. C., Vanden Eynden, E., Vandamme, A.-M., Deforche, K. & De Oliveira, T. (2018). Genome Detective: an automated system for virus identification from high-throughput sequencing data. Bioinformatics, 35 (5) 871–873.

Who. (2020). WHO Director-General’s opening remarks at the media briefing on COVID-19 [Online]. Available: https://www.who.int/emergencies/diseases/novel-coronavirus-2019/situation-reports [Accessed 26 July 2020].

Zhou, P., Yang, X. L., Wang, X. G., Hu, B., Zhang, L., Zhang, W., Si, H. R., Zhu, Y., Li, B., Huang, C. L., Chen, H. D., Chen, J., Luo, Y., Guo, H., Jiang, R. D., Liu, M. Q., Chen, Y., Shen, X. R., Wang, X., Zheng, X. S., Zhao, K., Chen, Q. J., Deng, F., Liu, L. L., Yan, B., Zhan, F. X., Wang, Y. Y., Xiao, G. F. & Shi, Z. L. (2020). A pneumonia outbreak associated with a new coronavirus of probable bat origin. Nature, 579 (7798) 270–273.

Zhu, N., Zhang, D., Wang, W., Li, X., Yang, B., Song, J., Zhao, X., Huang, B., Shi, W. & Lu, R. (2020). A novel coronavirus from patients with pneumonia in China, 2019. New England Journal of Medicine.

